# Mutation-Induced Effects on Rac1 Conformational Dynamics: Implications for Therapeutic Targeting

**DOI:** 10.1101/2025.09.04.673117

**Authors:** Busra Ozguney, Betul Uralcan, Saliha Ece Acuner, Turkan Haliloglu

## Abstract

Understanding the conformational dynamics of proteins, particularly small GTPases like Rac1, is vital for elucidating their functional mechanisms and developing targeted therapies. Rac1, pivotal in cellular processes, toggles between inactive GDP-bound and active GTP-bound states, regulated by guanine nucleotide exchange factors (GEFs) and GTPase-activating proteins (GAPs). Mutations, such as Rac1’s spontaneously activating oncogenic gain-of-function mutation P29S, associated with cancer, disrupt this equilibrium, leading to aberrant signaling. Traditional drug targeting of Rac1 is challenging due to its biological complexity and the lack of accessible active sites on its surface, necessitating alternative strategies. We propose a computational framework integrating Molecular Dynamics (MD) simulations and Elastic Network Models (ENM) to explore conformational dynamics. Our findings highlight the interplay between Mg^2+^ binding and conformational ensembles, revealing enhanced conformational heterogeneity in both inactive and active states upon P29S mutation. The critical location of P29S, Mg^2+^ coordination site, and GDP/GTP binding pocket with respect to global hinges provides mechanistic insight into how this mutation disrupts normal protein function through altered metal coordination dynamics. Furthermore, we identified strategic positions as potential “*rescue mutation*” sites, with T75A showing particular promise in mitigating the destabilizing conformational effects of P29S. Overall, this work provides insights into Rac1’s dynamic behavior and offers a foundation for targeted drug design strategies.

## Introduction

Proteins are essential for life to exist, and their dynamic nature allows them to visit various distinct functional states to act for certain biological function (1, 2). Each state consists of many populated substates that have similar structural and dynamic properties separated by low or high energy barriers (hills) yielding a multi-level hierarchy in evolutionary optimized Gibbs free energy landscape (3). Elucidating and accurately representing conformational substates that each state comprised is crucial for bridging protein dynamics and functionality and would yield novel strategies target challenging therapeutic proteins such as small GTPases that have limited accessible binding sites on their surface (4–6).

Ras-related C3 botulinum toxin substrate 1 (Rac1) is one of the small GTPases that has crucial roles in cell proliferation, adhesion, and migration functioning through switching between two globally distinct conformational states: inactive GDP-bound and active GTP-bound (7). In its active state, Rac1 interacts with downstream effectors such as PAK1 to involve signaling pathways by which Rac1 maintains its cellular functions (8–10). The ON/OFF state switch, i.e. GDP/GTP cycling, mechanism is tightly controlled by guanine nucleotide exchange factors (GEFs) which are responsible for replacing GDP with GTP by catalyzing GDP dissociation and GTPase-activating proteins (GAPs) enhancing the intrinsic hydrolysis activity to convert GTP to GDP (11–14).

Hot spot mutations of Rac1 are observed in many human cancers contributing cancer development, progression, and metastasis by causing abnormalities in the switch mechanism (15–17). In particular, gain-of-function P29S is associated with melanoma progression by increasing the effector activation (16–18). Upon this mutation, GDP/GTP nucleotide exchange rate increases, leading to spontaneous activation of the protein while maintaining the intrinsic GTP hydrolysis ability (18, 19). The increased transition rate was associated with the shift of the conformational equilibrium between the substates of the inactive Rac1 (20), but this hypothesis remains to be further explored since it gives limited explanation for the increased binding affinity towards downstream effectors in the active state upon P29S mutation (20). Moreover, P29S Rac1 engages in resistance mechanisms in melanoma by conferring resistance to RAF inhibitors, leading to increased cell viability and tumor growth despite targeted therapy (21).

Taken together, developing a protocol which aims to target P29S Rac1 has become attractive yet challenging task since Rac1 is not a classical druggable target (16–22), and it is impossible to create GTP-competitive Rac1 inhibitors (23).

Determining and manipulating the dynamics of substates that enables Rac1 to interact with its partners might be used as a novel strategy to rationally target Rac1 (23). Recently, it has been shown that Rac1.GDP exist in conformational equilibrium between at least two conformations with having different Mg^2+^ binding affinities depending on the ability of residue T35 coordination of Mg^2+^ (20). Moreover, according to many experimental and computational studies, active states of Rac1 (24–26) and Rac1 homologous proteins, such as Ras (27–36), RhoA (37, 38), Cdc42 (39), Rheb (40), and Ran (41), have at least two substates, state 1 and state 2, distinguished by the side chain conformations of either T35 or Y32 with respect to GTP yielding lack of or formation of hydrogen bond, respectively. Exploring the alleged substates in active state and understanding the driving force(s) that trigger conformational shifts between substates, would further elucidate the switch mechanism enabling control of the Rac1 interactions with its downstream and upstream effectors.

Towards this, we designed a computational scheme that combines information gathered from classical Molecular Dynamics (cMD) simulations, Replica Exchange Molecular Dynamics (REMD) simulations and Elastic Network Models (ENM) analysis. This integrated approach allows investigation of conformational ensemble shifts and dynamics underlying functional modifications in both inactive (GDP-bound) and active (GTP-bound) states of wild-type and P29S mutant Rac1. This framework reveals potential “rescue mutation” sites that, upon perturbation, can diminish the conformational and dynamic effects of the oncogenic P29S mutation. Analysis of these four distinct systems demonstrates a close connection between Mg^2+^ binding cavity status and populated conformational ensembles, ultimately revealing the conformational shifts and fast-cycling properties induced by the P29S mutation.

## Results

Rac1, a member of the Rho family of small GTPases, adopts a canonical Ras-like topology comprising a six-stranded β-sheet core surrounded by five α-helices. The 177-residue protein harbors a highly conserved nucleotide-binding pocket that exhibits high affinity for guanine nucleotides, accommodating GDP in the inactive state and GTP in the active conformation **(Figure 1)**. This pocket also binds to Mg^2+^ cofactor, essential for facilitating the guanine nucleotide exchange cycle by mediating GDP release and GTP hydrolysis. The Mg^2+^ coordination is mediated by the conserved residues T17 (p-loop), T35 (Switch 1), and D57 (Switch 2) **(Figure 1)**. Notably, the oncogenic P29S mutation, substituting a proline with serine at position 29 in the Switch I region, has been implicated in dysregulating the transition from the inactive GDP-bound to the active GTP-bound state. To elucidate the impact of this mutation on Rac1’s conformational landscape, we employed temperature-replica exchange molecular dynamics (REMD) simulations.

**Fig. 1.**
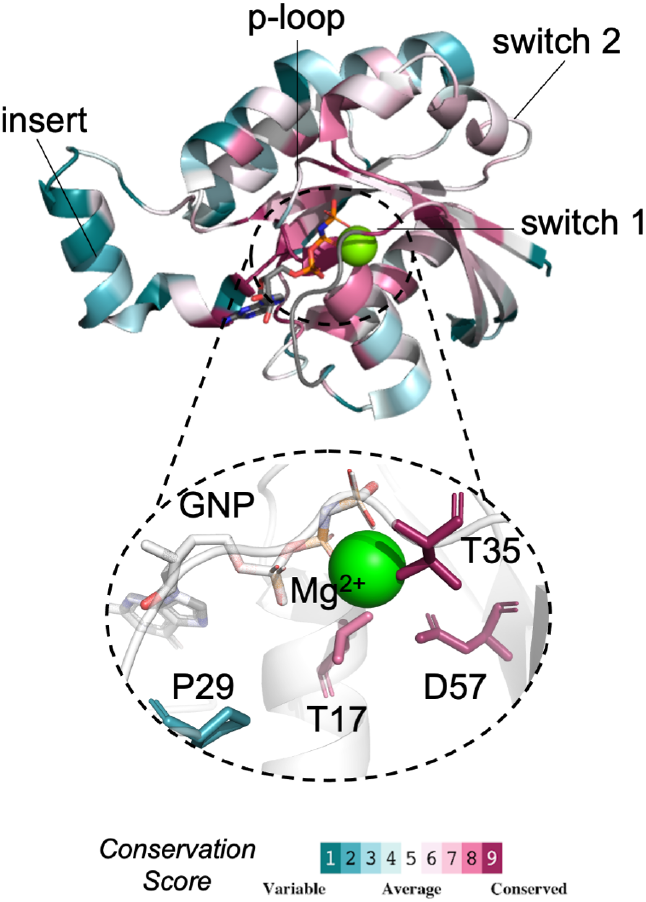
Structural and dynamic characterization of the Rac1 nucleotide-binding pocket. Crystal structure of wild-type Rac1 bound to the non-hydrolyzable GTP analog GNP (PDB ID: 3TH5), highlighting the nucleotide-binding pocket. The protein is color-coded by sequence conservation, emphasizing the highly conserved Switch I (N26-M45) and Switch II (A59-Q74) regions (red) crucial for nucleotide binding and conformational changes. The GDP/GTP substrate and Mg^2+^ cofactor (green sphere) are shown in the binding pocket.

### Inadequacy of Conventional Structural Descriptors in Representing Rac1 Dynamics Changes Due to P29S Mutation

First, we analyzed global variables, including the RMSD of C*α* atoms, R_*g*_ of C*α* atoms, and SASA, to understand structural changes caused by the P29S mutation in both inactive and active states. Results indicate that the inactive state, regardless of being wild-type or mutant, undergoes significant structural changes and samples more expanded conformations with increased solvent accessible surface area compared to the active state **(Figure S3)**. However, the global variables considered exhibit only slight changes in the mutant relative to the wild-type.

These findings suggest that conformational dynamics differ between the inactive and active states, with the former being more susceptible to structural changes. Nonetheless, the P29S mutation does not significantly alter the overall structure in either state, indicating changes in dynamics.

### P29S mutation expands Rac1’s conformational landscape and rewires inter-residue communication

To investigate the impact of the P29S mutation on Rac1’s conformational phase space in different states, we employed linear principal component analysis (PCA) on the Cartesian coordinates of Rac1’s inactive and active states obtained from Replica Exchange Molecular Dynamics (REMD) trajectories **(2A & C)**. PCA identifies dominant modes of motion with the largest variance, representing the most significant collective motions. Therefore, this approach overcomes the challenge of capturing directional changes in protein dynamics that scalar metrics like RMSD, R_*g*_ cannot capture.

The projection of coordinates onto the lower-dimensional subspace reveals that the P29S mutation induces enhanced conformational diversity in both inactive and active states. Specifically, the P29S mutant explores conformations inaccessible to wild-type along PC1 and PC2 in the inactive state, and along PC1 in the active state. The inactive state of wild-type Rac1 demonstrates greater conformational flexibility compared to its GTP-bound active state. These findings demonstrate that P29S alters Rac1’s energy landscape by inducing significant conformational heterogeneity rather than major structural alterations, with the inactive state being more susceptible to these dynamic changes.

We further investigated the collective motions represented by each principal component by analyzing conformations along PC1, PC2, and PC3 for each state. These collective motions were visualized Figure **(2B & D)** and quantified by calculating the RMSF values of C*α* atoms to identify the residues contributing most to the characterized motion **(Figure S4A & D)**. In the inactive state, PC1 of the P29S mutant describes increased separation between switch 1 and switch 2 regions, a motion represented by PC2 in wild-type Rac1 with less fluctuations in switch 2. The P29S mutation enhances mobility of residues in switch 1 and insert regions (D124-T135) along the first three principal components, suggesting a more collective motion modulating the opening/closing of the switch regions.

In the active state, the dominant motion (PC1) in both variants involves switch 1 opening, with P29S exhibiting enhanced mobility **(2D & S4D)**. Unlike the inactive state, switch 2 remains relatively stable in both variants. P29S shows collaborative motion between switch 1 and switch 2 in PC2, and coordinated movement of switch 1, β3 (K49-T58), and switch 2 in PC3, collectively resulting in favoring partial opening of the nucleotide binding pocket. In contrast, wild-type Rac1 shows increased switch 2 mobility in PC2 and N-terminal loop motions of switch 1 in PC3. These results demonstrate that the P29S mutation reshapes Rac1’s collective dynamics in a state-dependent manner, modifying switch mobility and altering the balance of nucleotide-pocket opening motions.

To gain additional insights into the correlated motions and allosteric communication underlying these conformational changes, we complemented our PCA with Dynamic Cross Correlation (DCCM) analysis and community network analysis **(Figure 3 & S5)**. The DCCM analysis revealed that P29S alters correlation patterns in both states **(Figure S5)**. In the inactive state, the mutation induces increased anti-correlated motions between switch 2 and the N-terminal, and between switch 2 and residues located between switch 2 and insert regions (T75-K123). Conversely, these residues (T75-K123) gain correlative motion with the N-terminal. In the active state, P29S causes loss of correlation between switch 1 and both switch 2 and residues T75-K123, as well as between the insert region and the C-terminal.

**Fig. 2.**
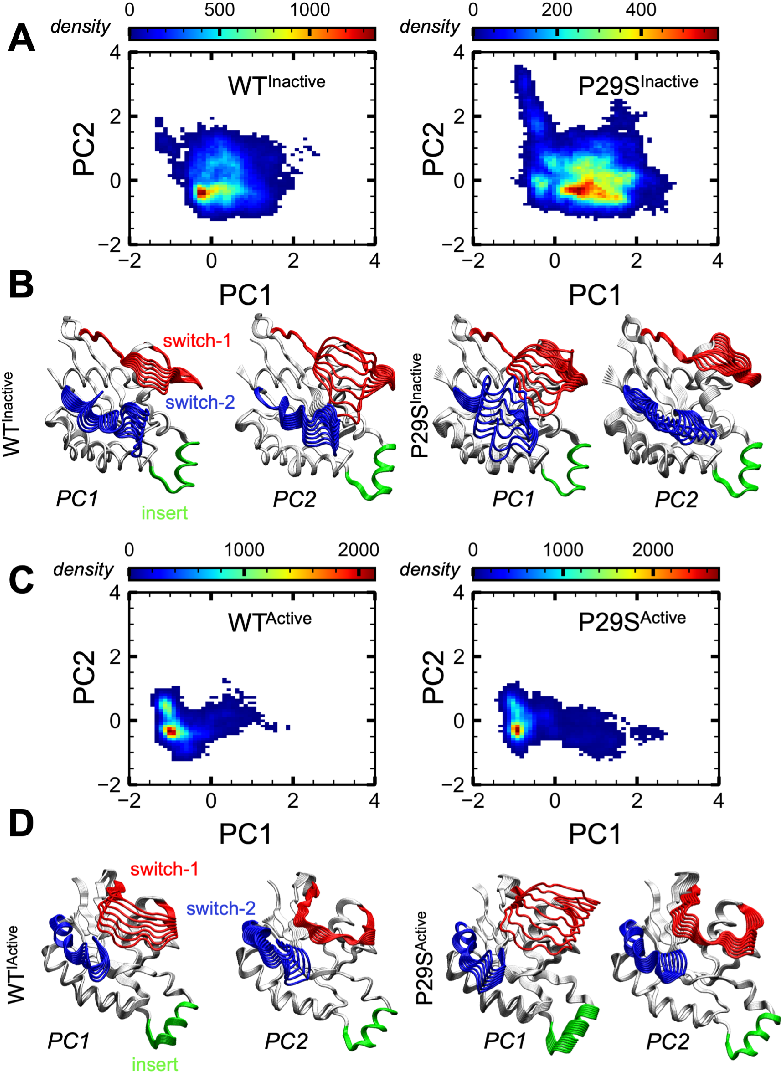
Conformational Phase Space Analysis of Rac1 Wild-Type and P29S Mutant Using Principal Component Analysis (PCA). **(A, C)** Density scatter plots of the first two principal components (PC1 and PC2) for Rac1 wild-type and P29S mutant in **(A)** inactive and **(C)** active states. These plots illustrate the conformational landscapes explored by each system, with PC1 and PC2 collectively accounting for approximately 30% of the total variance. The color gradient represents the density of conformations, with red indicating higher density. **(B, D)** Visualization of collective motions along PC1 and PC2 projected onto the X-ray structure of the Rac1 wild-type active state (PDB ID: 3TH5) for **(B)** inactive and **(D)** active ensembles. These projections illustrate the predominant structural fluctuations captured by PCA. The comparison between wild-type and P29S mutant in both states highlights the increased conformational heterogeneity induced by the P29S mutation. This analysis reveals distinct differences in the conformational ensembles sampled by the wild-type and mutant proteins, providing insights into the molecular basis of the mutation’s effects on Rac1 function.

**Fig. 3.**
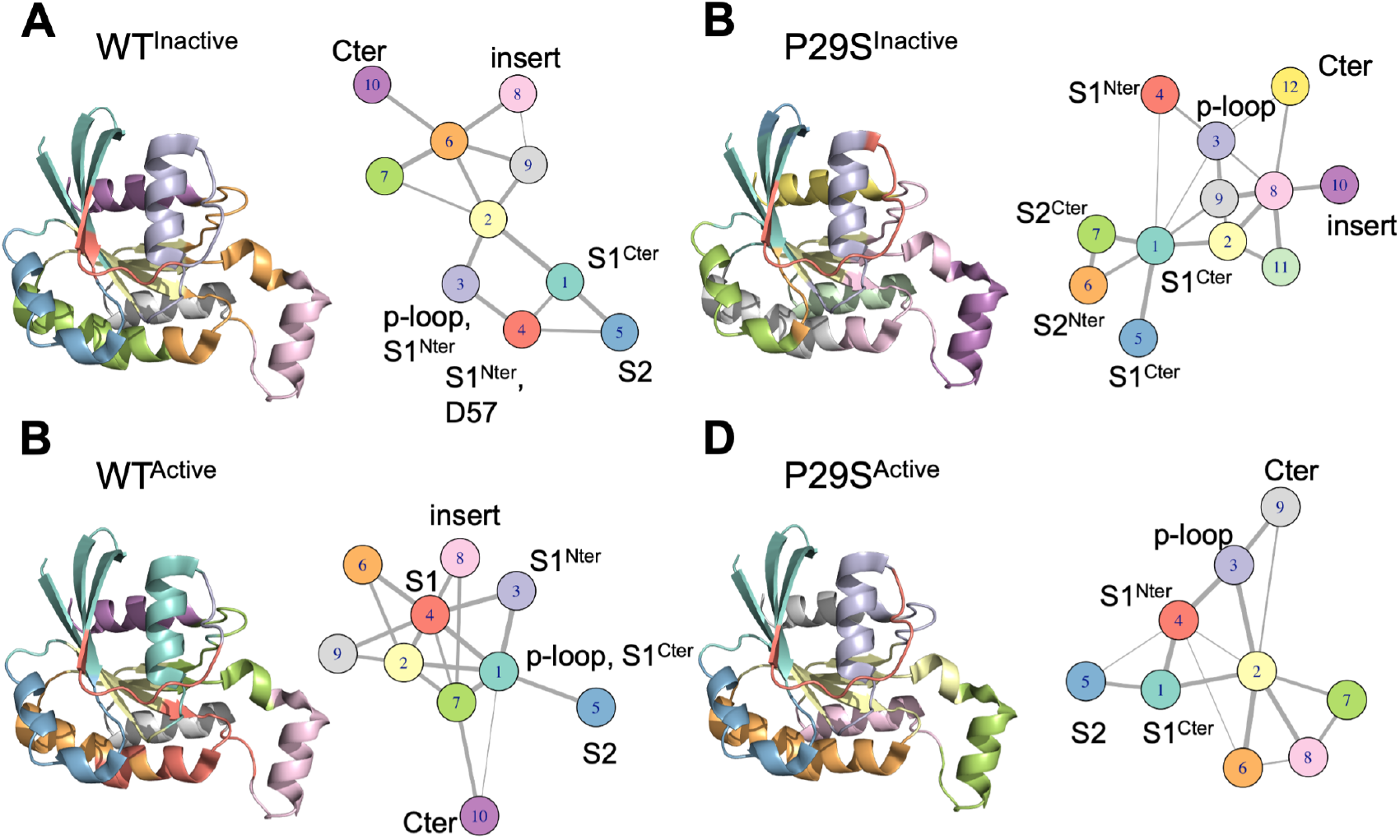
Community network analysis reveals P29S-induced alterations in Rac1’s allosteric communication. Community structures of Rac1 in **(A)** wild-type inactive, **(B)** P29S mutant inactive, **(C)** wild-type active, and **(D)** P29S mutant active states. Left: Communities mapped onto the Rac1 wild-type active state structure (PDB ID: 3TH5). Right: Corresponding 2D network representations. Distinct communities are color-coded consistently between 3D and 2D depictions. The P29S mutation induces notable reorganization of allosteric communities in both inactive and active states, particularly affecting switch regions and their interactions.

These findings are further supported by community network analysis, which identifies groups of highly negatively or positively correlated residues (communities) to reveal allosteric communication pathways within the protein ensemble. Our results revealed different numbers of communities in wild-type and mutant states **(Figure 3)**. Wild-type exhibits 10 communities in both states, while P29S shows 12 and 9 communities in inactive and active states, respectively. The increased fragmentation in the inactive state suggests P29S breaks larger coordinated regions into smaller units, while the decreased communities in the active state indicates that some previously separated regions now move in a more coordinated fashion.

Detailed examination of these communities reveals specific changes in inter-residue correlations consistent with the DCCM patterns described above. In the inactive state, switch 1 residues (I33-N39) and residues D57 and T58, which exist as a single community in the wild-type, are separated into different communities in the P29S mutant. The switch-2 region is divided into two communities, both losing correlation with the C-terminal of switch 1 (S41, N43-P50) **(Figure 3A-B & S5, Table S1&S2)**. In the active state, the P29S mutation expands a community including A27-G30 to encompass E31-D38, including key residues Y32 and T35. This community gains anti-correlation with the switch-2 region in the mutant **(Figure 3C-D & S5, Table S3&S4)**. Additionally, the p-loop becomes correlated with the C-terminal of the protein (S158-Q162, G164) and loses correlative motion with switch 2 in P29S. Collectively, these findings reveal a mutation-induced rewiring of Rac1’s allosteric network that manifests differently in the inactive and active states. While the inactive state changes primarily involve switch 2, the active state alterations are predominantly mediated through switch 1.

To further characterize these phenomena, we tracked the center of mass (CoM) distance between T35 (switch 1) and E62 (switch 2), revealing increased separation between switch regions in the P29S inactive state, indicated by a distinct shift towards larger values **(Figure S4C)**. While the inactive state is generally more flexible than the active state, RMSF profiles showed that P29S did not cause further increases in flexibility; instead, the p-loop, switch 1, and the insert region displayed trends toward slight stabilization relative to the inactive WT **(Figure S4B)**. These effects are consistent with nucleotide-pocket exposure without additional global destabilization. This trend is persisted but is less pronounced in the active state, in which switch 1 mobility is enhanced while preserving stability of switch 2 enabling partial opening of the nucleotide-binding pocket **(Figure S4E & F)**.

These detailed analyses demonstrate that local changes in inter-residue correlations and community structures drive the global conformational dynamics observed in PCA. The P29S mutation rewires Rac1’s internal communication networks, potentially impacting its interactions with binding partners and regulatory proteins. Many of the identified changes are centered around the nucleotide binding pocket. Given the central role of this region in Rac1 function and the observed perturbations in switch dynamics, further investigation into the local changes around the nucleotide binding pocket and the underlying mechanisms driving the differences in accessible conformational space for each state is needed.

### Conformational dynamics of T35 and Mg^2+^coordination in wild-type and P29S Rac1

While nucleotide exchange drives the transition between inactive and active states of small GTPases, the mechanisms underlying substate transitions within each state remain poorly understood. Previous studies on Rac1 homologs have utilized distances between key residues (Y32, T35) and the nucleotide or Mg^2+^ to characterize these substates (27, 37, 39). Building upon these findings, we investigated the conformational dynamics of wild-type and P29S Rac1 in both inactive and active states, focusing on the distances between the center of masses of T35 and Mg^2+^, Y32 and the nucleotide, and T35 and the nucleotide with the aim of elucidating the potential underlying driving force for the enhanced conformational heterogeneity observed in our PCA results for the mutant variant.

Probability distribution analysis revealed distinct conformational populations, particularly in T35-nucleotide and T35-Mg^2+^ distances (Figure S6). The Y32-nucleotide distance increased in the inactive states due to the absence of the γ-phosphate which is involved in hydrogen bonding with Y32, but remained largely unchanged in the active states. These observations suggest that T35 conformation plays a primary role in substate transitions, with Y32 conformation changing as a consequence. To elucidate the impact of the P29S mutation on Mg^2+^-coordinating residues (T17, T35, and D57), we examined their relative positioning **(Figure S7 & S8** and constructed free energy landscapes (FELs) using T35-Mg^2+^ and T17-D57 distances as order parameters for both inactive **(Figure 4A)** and active **(Figure 4B)** states.

**Fig. 4.**
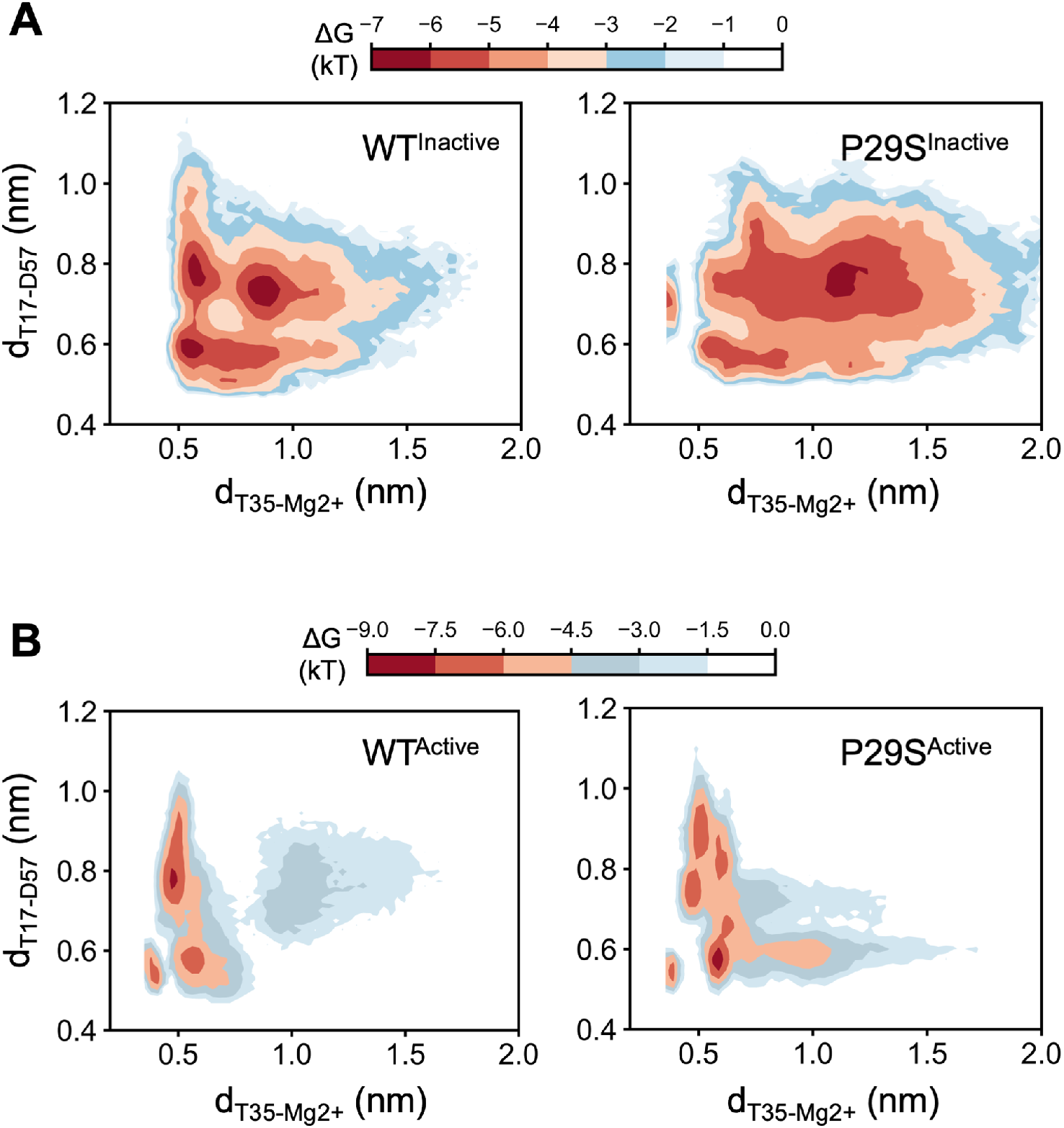
Free Energy Landscapes Reveal P29S-Induced Local Changes in Rac1’s Nucleotide Binding Cavity. Free energy landscapes (FELs) constructed using center of mass (CoM) distances between T35 and Mg^2+^, and between T17 and D57 as order parameters for (A) inactive and (B) active states of wild-type (left) and P29S mutant (right) Rac1. Color scale represents relative free energy (kT), with darker red indicating lower energy (more populated) states **(A)** In the inactive state, P29S mutation leads to a broader distribution of conformational states, particularly along the T35-Mg^2+^ axis, indicating increased flexibility in Mg^2+^ coordination. **(B)** The active state shows a more constrained landscape in wild-type Rac1, which becomes notably more heterogeneous upon P29S mutation, with the emergence of new energy minima. These FELs demonstrate increased conformational heterogeneity upon P29S mutation in both states. This altered energy landscape suggests a mechanism for the observed changes in nucleotide binding and exchange rates in the P29S mutant.

In the inactive state, wild-type Rac1 FEL revealed three populated basins with distinct T17-D57 conformations (close: 0.5-0.6 nm; distant: 0.7-1 nm) and two T35 conformations leading to either functional (0.5-0.7 nm) or impaired (>0.7 nm) Mg^2+^ coordination **(Figure 4A left panel)**. Distances of T17, T35, and D57 to Mg^2+^ showed bimodal distributions, allowing or disallowing coordination with almost equal probabilities **(Figure S7A)**. Considering T17-T35 and T35-D57 distances are always within the same range, and T17 and D57 are allowed to be positioned more flexible with respect to each other, T35 emerged as the most critical player in driving Mg^2+^ coordination. The P29S mutation increased conformational heterogeneity around the nucleotide binding cavity, enhancing configurations with impaired Mg^2+^ coordination by T35 and loss of T17-D57 interaction **(Figure 4A right panel)**. Distance distributions of Mg^2+^ with coordinating residues shifted towards larger values compared to wild-type, indicating compromised coordination **(Figure S7B)**. Analysis of nucleotide contacts within 4.5 Å revealed a slight rearrangement in GDP interactions within the switch 1 region upon mutation, with F28 and I33 losing contacts with GDP that may be attributed to accelerated dissociation rates **(Figure S9)**. Although π-π interaction between F28 and GDP is significantly loss in the P29S mutant, wild-type Rac1 also samples conformations that have loss this interaction. In summary, P29S mutation in the inactive state induces the conformational plasticity of the nucleotide binding cavity due to the impaired Mg^2+^ coordination.

In the active state, wild-type Rac1 exhibited a more rigid conformational space with three major basins sampling a narrow range of T35-Mg^2+^ distances (0.4-0.7 nm) and one less populated basin showing lack of ion coordination by T35 (>1 nm) **(Figure 4B left panel)**. T17-D57 distances in the major basins equally sampled close and distant configurations. The P29S mutation led to increased heterogeneity and access to hidden substates, enhancing the propensity to sample configurations with impaired T35-Mg^2+^ coordination **(Figure 4B right panel)**. A new basin emerged in the mutant, characterized by loss of T35-Mg^2+^ coordination (>0.7 nm) but stable T17-D57 interactions. Notably, while T35-Mg^2+^ coordination was mostly maintained in the wild-type, T17-Mg^2+^ and D57-Mg^2+^ distance distributions showed an increased probability of loss of ion coordination (Figure S8A). In the mutant, T17-Mg^2+^ coordination was never lost, and the majority of D57-Mg^2+^ distances remained small **(Figure S8B)**. This suggests that ion coordination in the mutant might be recovered more easily than in the wild-type after its loss. Investigation of protein-GTP interactions demonstrated an increased probability of loss of π-π interactions between F28-GTP and H-bonds between T35-GTP in the mutant, compensated by enhanced GTP interactions with S29-E31 **(Figure S9)**. The redistribution of contacts may contribute to the altered nucleotide binding and exchange properties observed in the P29S mutant. In contrast to inactive state, in the active state P29S strengths the Mg^2+^ coordination by T17 and D57, and, thereby, allowing nucleotide binding cavity to change between configurations without losing the ion coordination completely.

In conclusion, our analysis reveals that Rac1 adopts different interconvertible conformational substates within both inactive and active states, with Mg^2+^ coordination playing a crucial role in these transitions. The P29S mutation induces conformational plasticity in the nucleotide binding cavity, leading to altered Mg^2+^ coordination dynamics and nucleotide interactions. These changes reduce energy barriers between substates in inactive and active states, resulting in a conformational shift in the energy landscape that likely contributes to the fast-cycling property of the mutant. Our findings provide mechanistic insights into the enhanced nucleotide exchange rates and altered signaling properties observed in the P29S Rac1 mutant, offering new insights into the molecular mechanisms underlying Rac1-associated pathologies and potentially informing future therapeutic strategies.

### Identification and characterization of rescue mutations stabilizing P29S Rac1 active state

Having established the profound impact of the P29S mutation on Rac1’s conformational landscape and Mg^2+^ coordination, we sought to identify compensatory mutations that could mitigate these effects, particularly in the active state. As the functionally relevant state for effector binding, perturbations in the active state directly impact Rac1’s biological activity. Moreover, the more localized conformational changes in the active state (primarily switch 1 mobility) suggest that targeted stabilization strategies might be more achievable than addressing the extensive perturbations observed in the inactive state. Our primary objective was to discover mutations capable of reducing the P29S-induced conformational heterogeneity and stabilizing the switch 1 region, potentially restoring normal Rac1 function and providing insights into therapeutic strategies.

While our initial PCA analysis elucidated global conformational changes induced by the P29S mutation, it was crucial to identify the specific residues driving these large-scale motions. To this end, we employed Gaussian Network Model (GNM) analysis, a complementary approach to PCA that offers a distinct perspective on protein dynamics (42, 43). GNM utilizes a coarse-grained approach, modeling the protein as a network of interconnected nodes represented by C*α* atoms. This method excels at identifying hinge residues, critical mechanical points in the protein structure that facilitate large-scale conformational changes.

In GNM analysis, the low-frequency modes often correspond to functionally relevant motions (24, 44–46). By focusing on the first two GNM modes, we aimed to capture the most significant collective motions of Rac1 and pinpoint the critical residues governing these motions. Notably, the P29S mutation site is in direct contact with a hinge residue (L18) of the slowest mode **(Table S5)**, which motivated the search for alternative hinge residues to compensate for the deformation caused by P29S.

Leveraging these insights, we implemented a systematic computational mutagenesis approach, targeting the hinge residues identified through GNM analysis of Rac1’s first two dynamic modes in the active state **(Figure S10 & Table S5)**. We hypothesized that perturbing hinge movements may trigger allosteric regulation of protein function by causing a population shift in the conformational ensemble (46). This approach could potentially reveal cryptic allosteric binding sites that do not compete with nucleotide binding and could modify the ensemble of conformations towards the wild type.

We generated an in-silico library of Rac1 variants by systematically mutating each identified hinge residue. Each variant, along with the wild-type and P29S mutant, underwent a series of conventional MD (cMD) simulations at progressively increasing timescales (10 ns, 50 ns, and 100 ns) with multiple replicas. This comprehensive approach allowed us to assess the impact of each mutation, the tendency on conformational heterogeneity and switch 1 stability in the active state. Our analysis encompassed evaluations of structural stability, residue correlations, and both intra-protein and protein-nucleotide/Mg^2+^ contact propensities around the nucleotide binding cavity **(unpublished data)**.

This rigorous screening process enabled us to identify two promising mutations that, when introduced alongside P29S, showed potential to counteract its destabilizing effects. To validate and characterize these double mutants, P29S+L19A and P29S+T75A, we subjected them to REMD simulations **(Figure 5A-B, & S10)**.

**Fig. 5.**
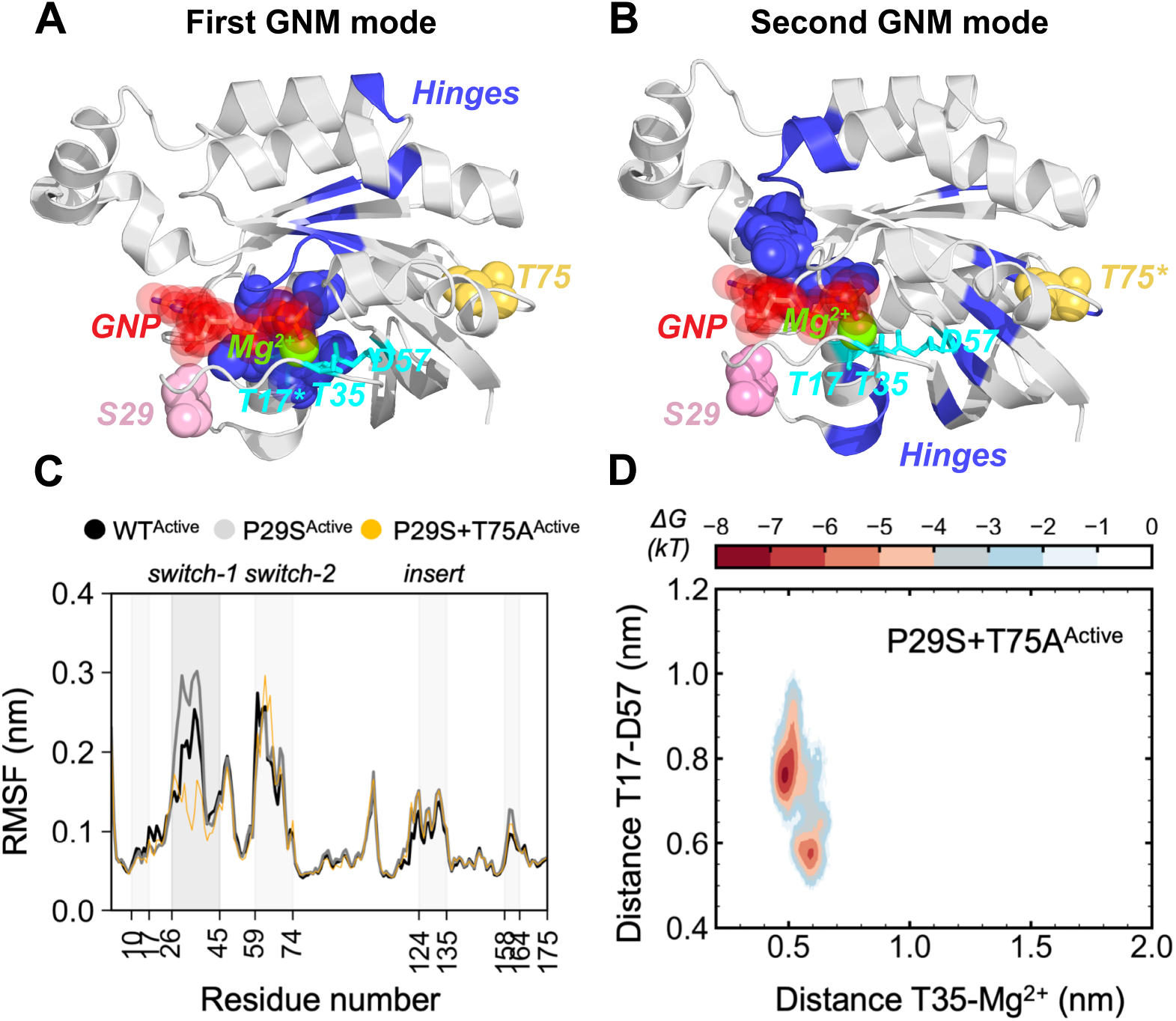
Identification and characterization of compensatory mutations in the active state of Rac1 using Gaussian Network Model (GNM) analysis and enhanced sampling techniques. **(A, B)** Visualization of hinge residues identified from the **(A)** first and **(B)** second GNM modes of Rac1. Hinge residues, highlighted in blue, represent flexible joints connecting rigid units and mobile loops within the protein structure. These residues were determined by analyzing minima in the mean-square fluctuation curves or correlation alterations, derived from the eigenvectors of the slowest GNM modes. Domain dissection through hinge resides could be seen further in Figure S10. **(C)** Root mean square fluctuation (RMSF) analysis of C*α* atoms in wild-type, P29S, and P29S+T75A Rac1 active states. The plot highlights the stabilizing effect of the T75A rescue mutation on the switch-1 region of P29S Rac1. **(D)** Free energy landscape (FEL) of the P29S+T75A rescue mutant, constructed using the distances between T35-Mg^2+^ and T17-D57 as order parameters. The FEL demonstrates reduced conformational plasticity of the nucleotide binding cavity in the rescue mutant compared to P29S Rac1. This analysis pipeline, combining GNM-guided mutagenesis with enhanced sampling techniques, enabled the identification and validation of potential compensatory mutations to mitigate the effects of the oncogenic P29S mutation in Rac1.

The REMD simulations revealed that the P29S+T75A variant exhibited significantly enhanced stability in the switch-1 region compared to both the wild-type and the P29S mutant in active state **(Figure 5C)**. Analysis of the conformational space sampled by the T35-Mg^2+^ distance and the T17-D57 distance showed a decreased propensity for conformations with impaired Mg^2+^ coordination in the T75A variant **(Figure 5D)**. These computational results suggest that the T75A mutation, when combined with P29S, may enhance the stability and functional integrity of the protein. However, it should be noted that these findings are based on in silico simulations and require further experimental validation to confirm their biological relevance.

## Conclusions

Over-activation of the small GTPase Rac1 due to mutations has been linked to various cancers [17]. Among these recurrent somatic missense mutations, P29S stands out as the most prevalent cancer-associated mutation in the Rho family GTPases and ranks as the third most common mutation observed in melanoma (18, 22, 47, 48). Despite extensive research on the P29S mutation (18–20, 24, 25), its rapid cycling behavior between GDP- and GTP-bound states remains unclear. In this study, we aimed to elucidate the conformational dynamics of the P29S mutation in both inactive and active states. Our focus was on the conformational heterogeneity and the mechanistic impact of the mutation on collective motions, utilizing enhanced sampling techniques. Additionally, we identified a potential “rescue mutation” site that could mitigate the destabilizing effects of the P29S mutation.

Our present study of Rac1’s conformational dynamics reveals a fundamental remodeling of protein’s energy landscape by the P29S mutation. In both inactive GDP-bound and active GTP-bound states, the mutant explores previously inaccessible conformational space exhibiting increased structural plasticity that manifests in enhanced nucleotide exchange rates and altered signaling properties observed in P29S Rac1. This enhanced conformational plasticity provides insights into the fast-cycling property of the P29S mutant, as the mutation allows the protein to populate different substates, some of which are hidden in the wild-type.

Through integration of linear dimensionality reduction with correlation analysis, we uncovered a mutation-induced rewiring of Rac1’s allosteric network that differs between inactive and active states. In the inactive GDP-bound state, the mutation triggers increased mobility of both switch regions, leading to enhanced exposure of the nucleotide-binding cavity. This structural perturbation likely facilitates GDP release, explaining the accelerated nucleotide exchange observed in previous biochemical studies (19, 20). In contrast, the active GTP-bound state shows selective enhancement of switch 1 mobility while maintaining switch 2 stability, resulting in a precisely controlled partial opening of the nucleotide-binding pocket that may optimize interactions with effector proteins.

Central to these conformational changes is the altered co-ordination of Mg^2+^, which emerges as a critical modulator of Rac1’s dynamics. In the inactive state GDP-bound state, P29S disrupts both T35-Mg^2+^ coordination and the critical T17-D57 interaction, destabilizing the nucleotide-binding pocket. The active GTP-bound state reveals a more nuanced mechanism: while T35-Mg^2+^ coordination becomes impaired, coordination by T17 and D57 strengthens, suggesting a compensatory mechanism that maintains partial nucleotide pocket stability. This state-specific modulation of metal co-ordination provides a structural framework for understanding the fast-cycling behavior of P29S Rac1.

Our analysis further reveals that the P29S mutation fundamentally alters Rac1’s conformational landscape in both inactive and active states. In the inactive state, the mutation shifts the conformational ensemble towards substates with impaired Mg^2+^ coordination, particularly by T35, and loss of the T17-D57 interaction. This alteration in Mg^2+^ binding affinity aligns with previous suggestions (19, 20) and may contribute to increased intrinsic GDP dissociation as well as GEF-induced GDP dissociation, explaining the decreased probability of the protein remaining in the inactive state. In the active state, while the P29S mutation enhances the propensity for configurations with impaired T35-Mg^2+^ co-ordination, it strengthens coordination by T17 and D57. This state-specific alteration in Mg^2+^ coordination dynamics, coupled with the increased conformational diversity, enables P29S.GTP to more readily adopt conformations suitable for binding downstream and upstream effectors.

The significance of metal ion coordination in protein function is further supported by recent work from Nam et al. (2024)(49), who demonstrated that proper Mg^2+^ coordination in adenylate kinase reduces non-functional conformational states and enhances catalytic activity (49). Similarly, our findings reveal that P29S mutation increases nucleotide-binding cavity plasticity through subtle perturbations in metal coordination, rather than through large conformational changes, thereby modulating the dynamics of functional regions such as switch 1 and switch 2. This mechanistic understanding of how metal coordination influences protein function suggests that targeting the stability of metal-coordinating residues could provide a novel therapeutic strategy.

Building upon these mechanistic insights, we employed an innovative approach combining GNM analysis with systematic computational mutagenesis to identify potential compensatory mutations. This strategy bridges the gap between global conformational changes and specific residues that could be targeted to modulate these changes. Our rigorous screening process revealed that the P29S+T75A double mutant exhibits significantly enhanced stability in the switch-1 region compared to both wild-type and P29S Rac1. Moreover, this variant showed a decreased propensity for conformations with impaired Mg^2+^ coordination, suggesting a potential restoration of normal Rac1 function. The identification of these compensatory mutations, particularly T75A, opens new avenues for therapeutic intervention by potentially counteracting the destabilizing effects of P29S.

Our findings provide a comprehensive framework for understanding the molecular mechanisms underlying Rac1-associated pathologies and offer a foundation for developing targeted therapeutic strategies. However, it is crucial to emphasize that these findings are based on in silico simulations and require further experimental validation. Future studies should focus on experimental validation of the predicted effects of T75A and other potential compensatory mutations on Rac1 stability, nucleotide binding, and effector interactions. Additionally, investigation of the functional consequences of these mutations in cellular contexts, particularly their impact on Rac1-mediated signaling pathways, will be essential. Further exploration of small molecule modulators that could mimic the stabilizing effects of compensatory mutations may lead to novel therapeutic agents.

In conclusion, our study provides unprecedented insights into how cancer-associated mutations can reprogram protein dynamics through subtle perturbations of metal coordination networks. By elucidating the intricate interplay between local structural changes and global functional outcomes, we have not only advanced our understanding of Rac1 biology but also paved the way for innovative therapeutic strategies targeting this crucial signaling protein in cancer and other diseases. These findings underscore the power of integrating advanced computational methods to unravel complex protein dynamics and identify potential therapeutic targets in challenging systems like small GTPases.

## Materials and Methods

Temperature-Replica Exchange Molecular Dynamics (REMD) simulations were conducted at 310 K and 1 bar, in which protein is modeled with Charmm36 force field along with TIP3P water model. **Structural Analysis:** All structural analysis were conducted using GROMACS (50). These included calculations of root mean square deviation (RMSD), root mean square fluctuation (RMSF), solvent accessible surface area (SASA), inter-atomic distances, and covariance matrices. **Principle Component Analysis (PCA):** PCA was performed on the C*α* atoms of the trajectories after concatenating all sampled frames from all constructs. The first three principal components (PCs) were extracted, and RMSF along these PCs were calculated using individual trajectories of each construct. This analysis allowed for the identification of dominant collective motions and their contributions to overall protein dynamics. **Free Energy Landscape Construction:** Two-dimensional free energy landscapes were constructed using the equation:

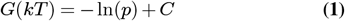

where *k* is the Boltzmann constant, *T* is temperature, *p* is the joint probability distribution, and *C* is an additive constant. The order parameters used were the center of mass distances between residue T35 and the Mg^2+^ ion, and between residues T17 and D57. This approach enables visualization of the conformational space explored by the protein and identification of energetically favorable states. **Protein-Nucleotide Contact Analysis:** Protein-nucleotide contacts were quantified using MDAnalysis (51, 52). Contacts were defined as heavy atoms of protein residues within 0.45 nm of heavy atoms of the nucleotide (GDP or GTP). This analysis provides insights into the specific interactions mediating protein-nucleotide binding. **Dynamic Cross-Correlation and Community Network Analysis:** Dynamic cross-correlation matrices (DCCM) were calculated by normalizing the covariance matrices of the trajectories for inactive and active states of wild-type Rac1 and the P29S mutant. The covariance of linear atomic displacements of C*α* atoms for residues i and j was calculated, with the resulting matrix normalized to have a diagonal of 1. Community network analysis was performed using Bio3D (53). A weighted network graph was constructed where nodes represent individual residues (based on C*α* atoms), and edge weights between nodes i and j represent their respective cross-correlation values. Network edges were added for residue pairs with cross-correlation values ≥ |0.4|, with edge weights calculated as - log(cross-correlation). Communities of highly correlated residues were identified using the edge-betweenness clustering algorithm developed by Girvan and Newman. This analysis provides insights into allosteric communication pathways within the protein structure. **Identification of Hinge Residues:** Hinge residues were identified using HingeProt (54). In this, we used crystal structure of P29S active state (PDB id: 3SBD) and divided protein structure into rigid parts and the hinge regions connecting them.

