## Supplementary Information for "Mutation-Induced Effects on Rac1 Conformational Dynamics: Implications for Therapeutic Targeting"

**Table S1. Residues belonging to each community revealed by community network analysis for wild-type Rac1 inactive state**

| Wild-type Rac1 inactive | |
|---|---|
| Community (size) | Residue number |
| 1 (26) | 2-8; 40-56; 75-76 |
| 2 (19) | 9-10; 77-82; 108-114; 152-155 |
| 3 (22) | 11-32 |
| 4 (9) | 33-39; 57-58 |
| 5 (16) | 59-74 |
| 6 (25) | 83-90; 115-121; 136; 156-164 |
| 7 (17) | 91-107 |
| 8 (14) | 122-135 |
| 9 (15) | 137-151 |
| 10 (13) | 165-177 |

**Table S2. Residues belonging to each community revealed by community network analysis for P29S Rac1 inactive state**

| P29S Rac1 inactive | |
|---|---|
| Community (size) | Residue number |
| 1 (22) | 2-8; 40-41; 43; 52-57; 75-78; 108-109 |
| 2 (12) | 9; 79-81; 110-113; 152-155 |
| 3 (16) | 10-25 |
| 4 (14) | 26-39 |
| 5 (9) | 42; 44-51 |
| 6 (4) | 58-61 |
| 7 (13) | 62-74 |
| 8 (28) | 82-91; 114-120; 135-137; 156-163 |
| 9 (16) | 92-107 |
| 10 (14) | 121-134 |
| 11 (14) | 138-151 |
| 12 (14) | 164-177 |

**Table S3. Residues belonging to each community revealed by community network analysis for wild-type Rac1 active state**

| Wild-type Rac1 active | |
|---|---|
| Community (size) | Residue number |
| 1 (43) | 2-7; 13-26; 39-57; 74-76; 159 |

| | |
|---|---|
| 2 (18) | 8-12; 77-81; 109-113; 152-154 |
| 3 (4) | 27-30 |
| 4 (25) | 31-38; 82-91; 93-96; 135-137 |
| 5 (16) | 58-73 |
| 6 (13) | 92; 97-108 |
| 7 (16) | 114-120; 155-158; 160-164 |
| 8 (14) | 121-134 |
| 9 (14) | 138-151 |
| 10 (13) | 165-177 |

**Table S4. Residues belonging to each community revealed by community network analysis for P29S Rac1 active state**

| Wild-type Rac1 active | |
|---|---|
| Community (size) | Residue number |
| 1 (30) | 2-7; 39-58; 74-77 |
| 2 (31) | 8-10; 78-85; 108-120; 151-157 |
| 3 (23) | 11-26; 158-163; 165 |
| 4 (12) | 27-38 |
| 5 (15) | 59-73 |
| 6 (22) | 86-107 |
| 7 (15) | 121-135 |
| 8 (15) | 136-150 |
| 9 (13) | 164; 166-177 |

**Table S5. Hinge residues depicted by GNM analysis**

| | |
|---|---|
| *First Slowest Mode* | 9, 10, 11, 12, 13 |
| *First Slowest Mode* | 15, 16, 17, 18, 19, 20 |
| *First Slowest Mode* | 80, 81 |
| *First Slowest Mode* | 97,98 |
| *First Slowest Mode* | 112, 113 |
| *First Slowest Mode* | 148, 149 |
| *First Slowest Mode* | 153, 154 |
| *First Slowest Mode* | 165, 166 |
| *Second Slowest Mode* | 6, 7 |
| *Second Slowest Mode* | 23, 24, 25 |
| *Second Slowest Mode* | 39, 40 |
| *Second Slowest Mode* | 55, 56 |
| *Second Slowest Mode* | 74, 75 |

| | |
|---|---|
| *Second Slowest Mode* | 77, 78 |
| *Second Slowest Mode* | 84, 85, 86, 87, 88, 89 |
| *Second Slowest Mode* | 116, 117 |
| *Second Slowest Mode* | 138, 139 |
| *Second Slowest Mode* | 151, 152, 153 |
| *Second Slowest Mode* | 163, 164 |

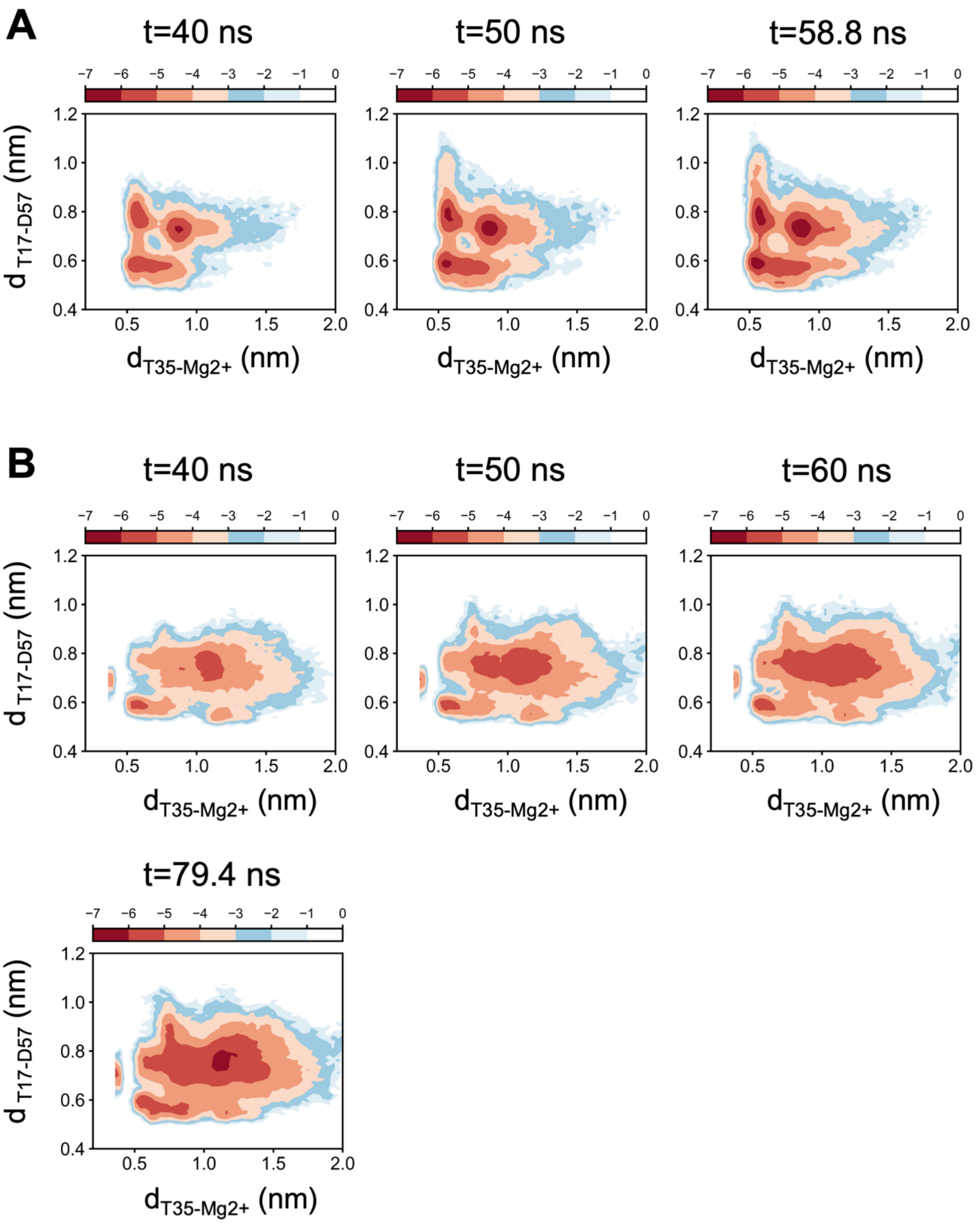

**Figure S1. Convergence of the REMD trajectories.** Free energy landscapes are constructed using the order parameters discussed in the main text for different intervals of the time space at 310 K for A) WT-inactive and B) P29S-inactive states. Results indicate the convergence of the trajectories as with increasing time, configurational space that is explored is not changing.

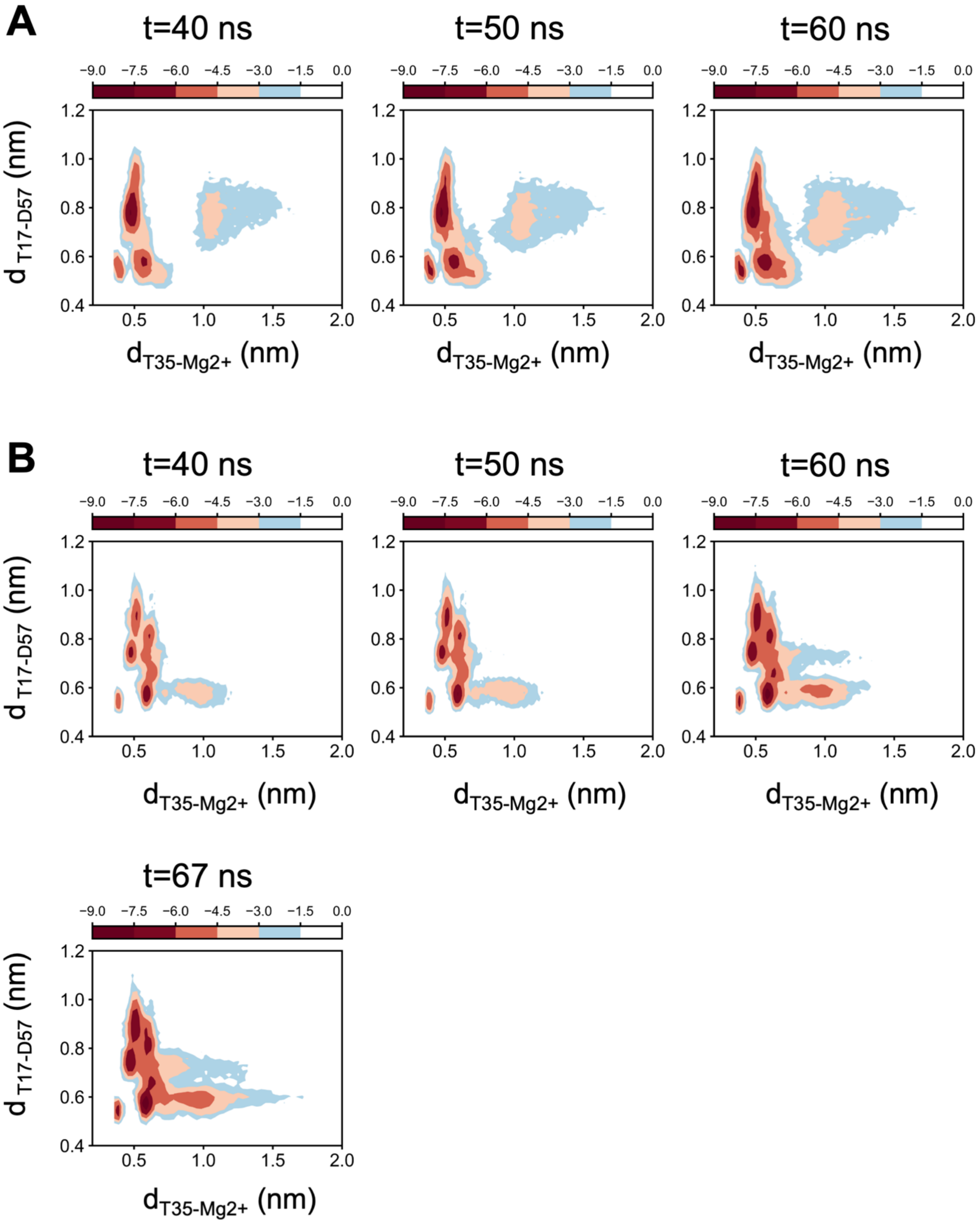

**Figure S2. Convergence of the REMD trajectories.** Free energy landscapes are constructed using the order parameters discussed in the main text for different intervals of the time space at 310 K for A) WT-active and B) P29S-active states. Results indicate the convergence of the trajectories as with increasing time, configurational space that is explored is not changing.

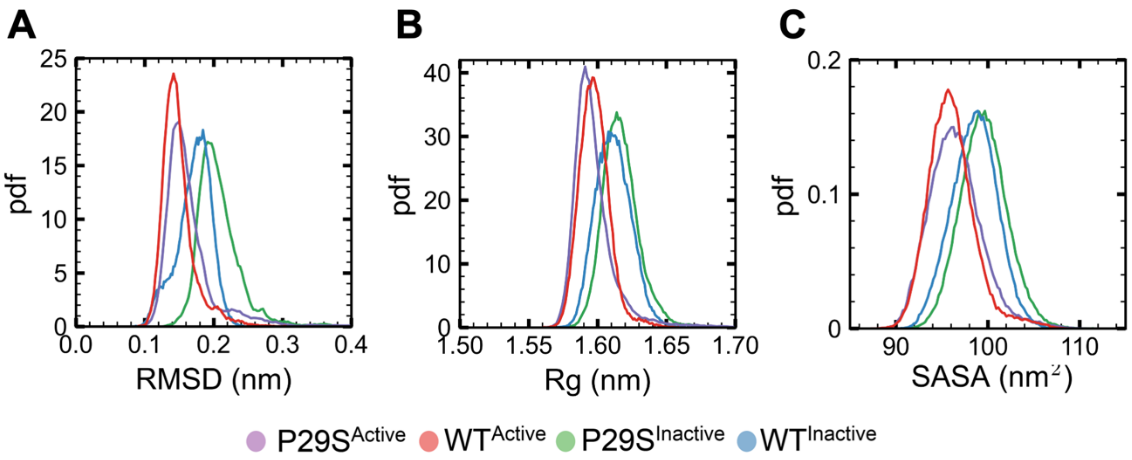

**Figure S3. Global variables are inadequate to characterize the conformational heterogeneity of Rac1.** A) Root mean square deviation (RMSD) of Ca atoms, B) Radius of gyration (Rg) of Ca atoms, and C) Solvent accessible surface area (SASA) of protein for wild-type Rac1 and P29S mutant in both inactive and active states. Results indicate global variables slightly change when compared inactive state to active state but are not able to capture the impact of the oncogenic P29S mutation.

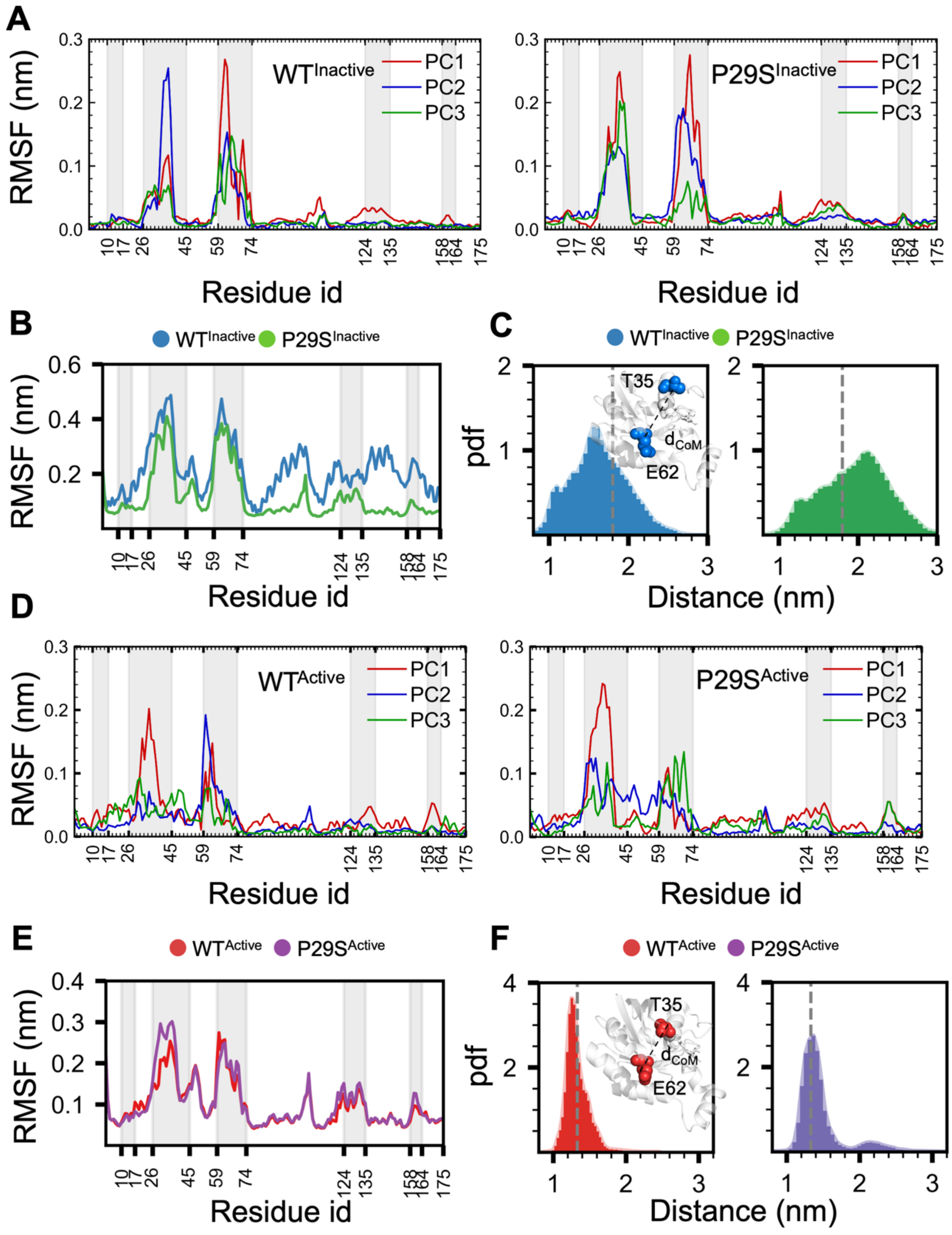

**Figure S4. Residue mobilities associated with principal components.** A) For inactive state, B) For active state.

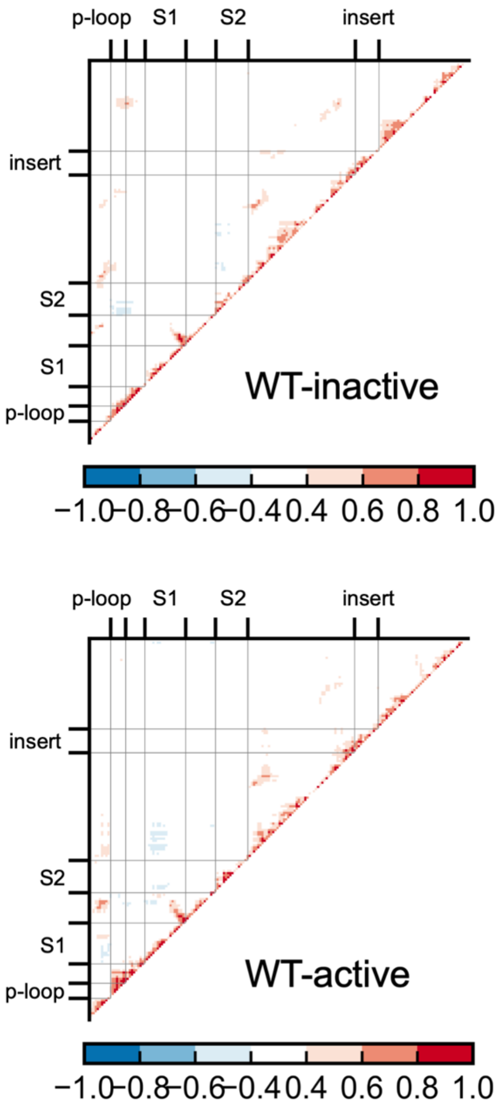

**Figure S5. Dynamic cross correlation maps of wild type and P29S mutant in inactive and active states.**

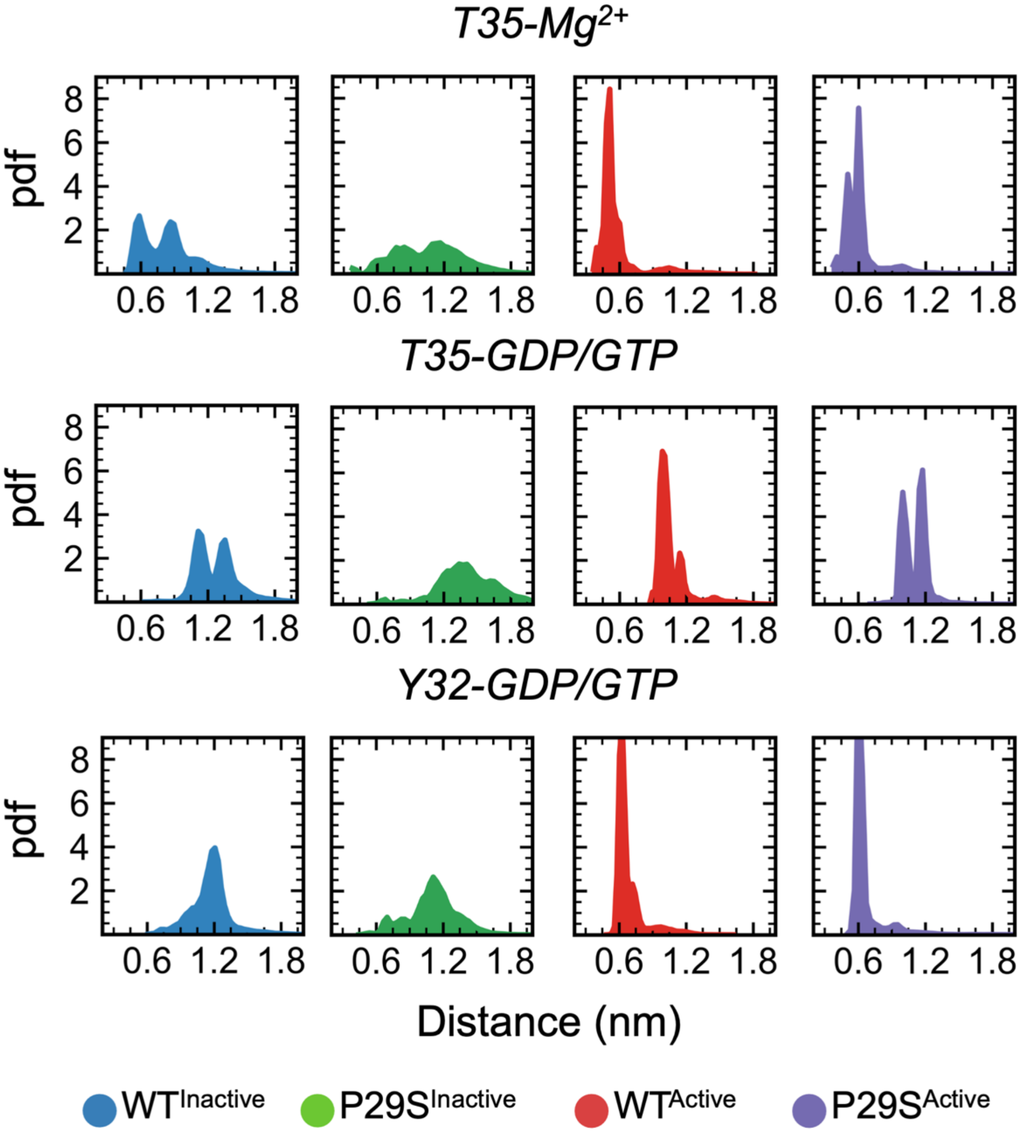

**Figure S6. Distance distributions of key interactions: T35-Mg2+, T35-nucleotide, and Y32-nucleotide.** These order parameters, known to characterize binding pocket promiscuity in Rac1 homologs (e.g., Cdc42, Raf, Rho), reveal conformational heterogeneity within the nucleotide-binding site.

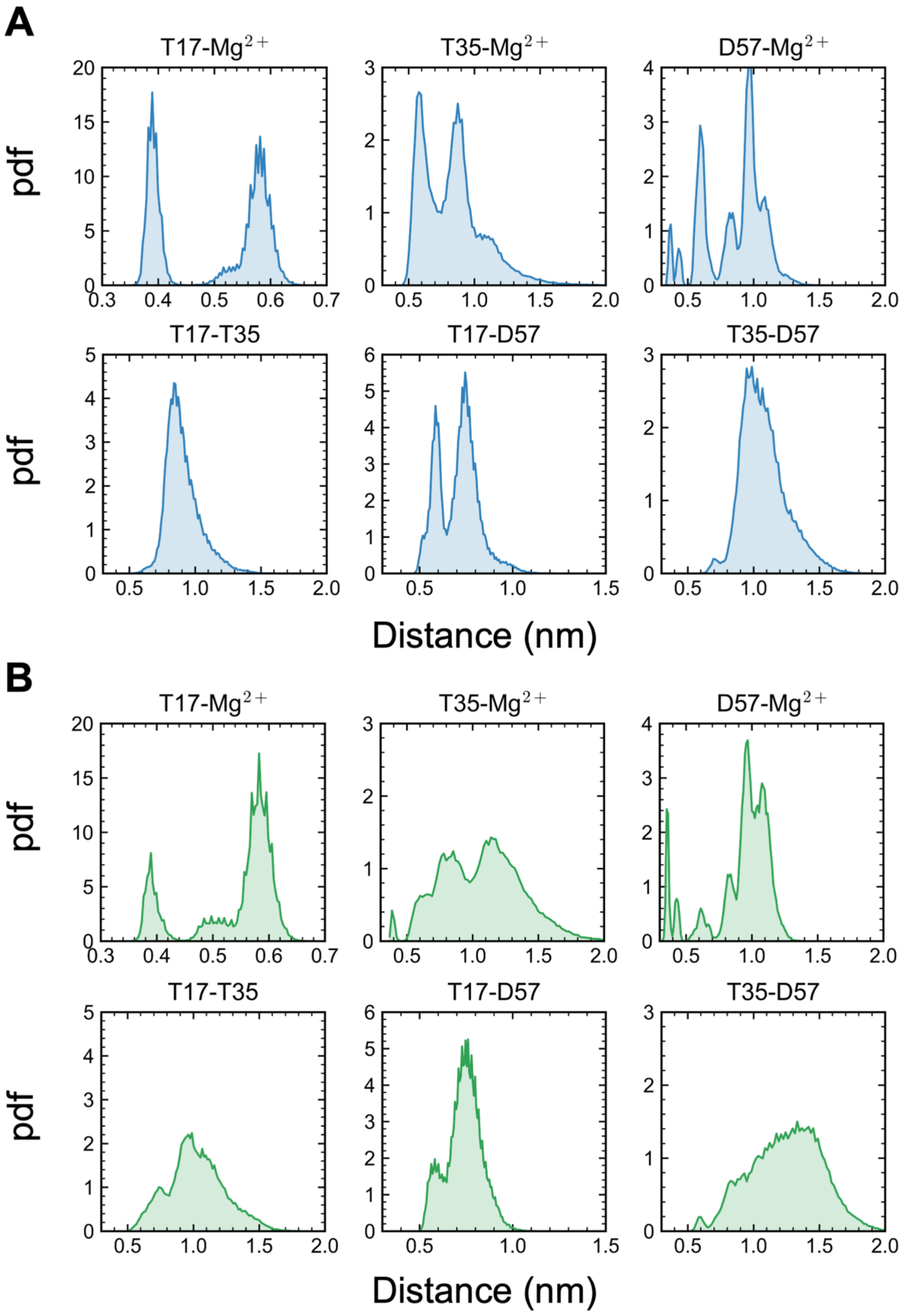

**Figure S7. Distance distributions of Mg2+ coordinating residues with respect to each other and ion in inactive state.**

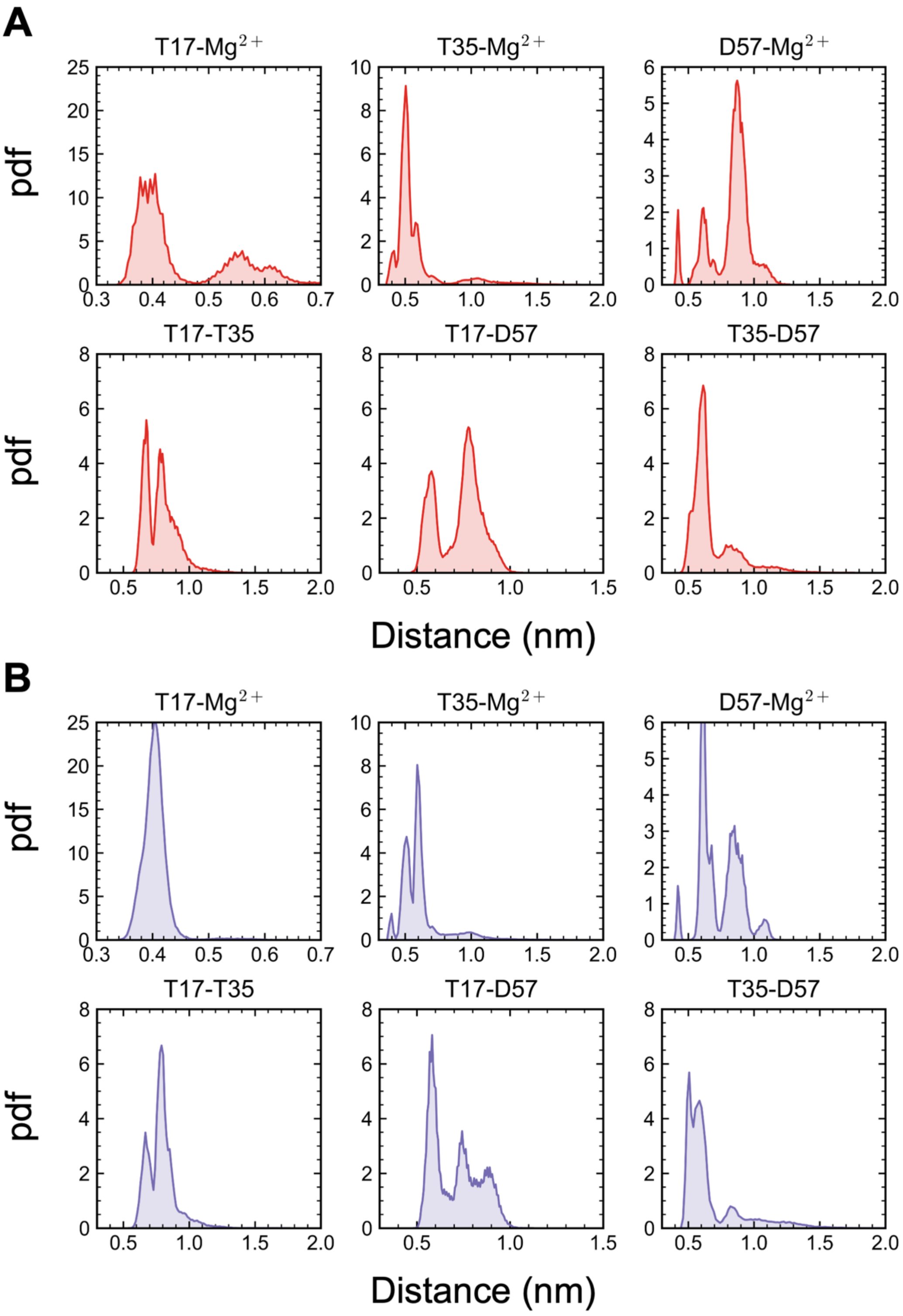

**Figure S8. Distance distributions of Mg2+ coordinating residues with respect to each other and ion in active state.**

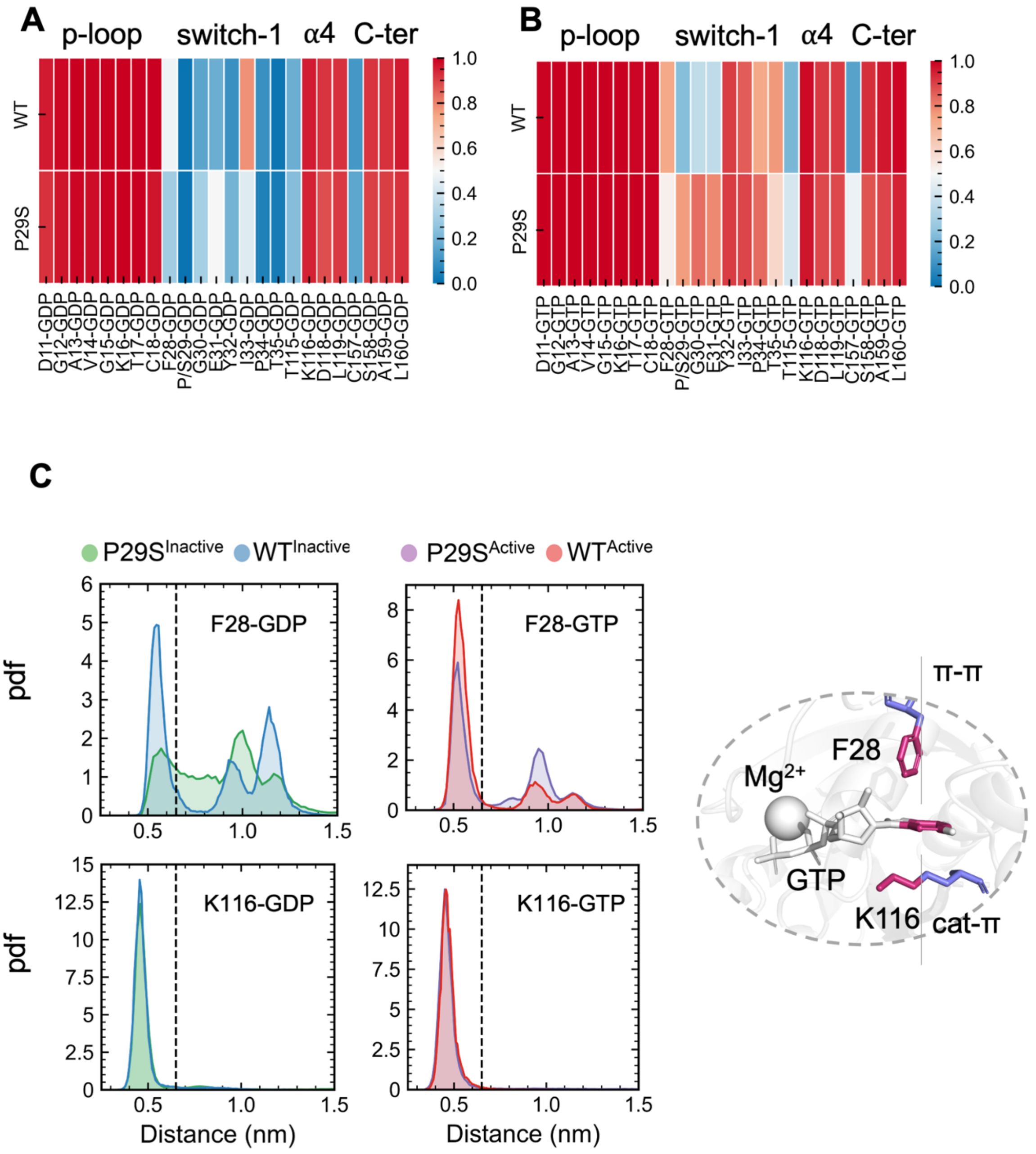

**Figure S9. Changes occur upon mutation in protein-nucleotide interactions (A)** Protein-GDP interactions within 0.45 nm in wild type and P29S mutant **(B)** Protein-GTP interactions within 0.45 nm in wild type and P29S mutant (C) Tracking stacking interactions stabilizing nucleotide in wild type through stacking sandwich. Nucleotide has a pi-pi stacking with aromatic ring of F28 and cat-pi stacking with K116. Center of mass distance of aromatic rings (for F28-nucleotide) and of aromatic ring to positively charged group of Lys (K116-nucleotide) distributions show Lys-nucleotide interactions are intact while Phe-nucleotide interactions are impaired upon mutation.

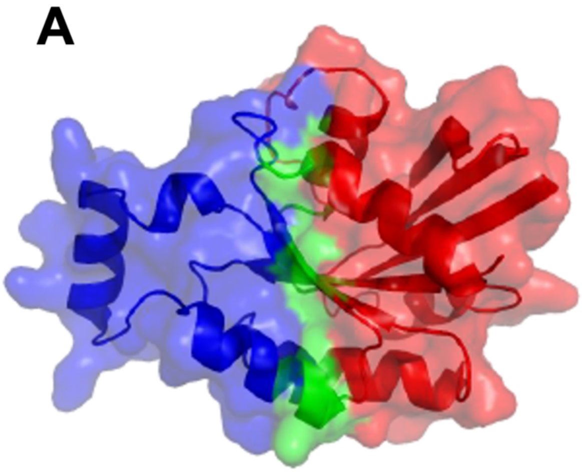

**Figure S10. Identification of Hinge Residues in P29S Rac1 in Active State Using Gaussian Network Model (GNM) Analysis (A)** First (lowest frequency) mode and **(B)** Second lowest frequency mode of Rac1 dynamics as predicted by GNM. Hinge residues, identified as local minima in the mode shapes, are depicted as green spheres on the Rac1 structure (PDB id: 3SBD). These residues represent key mechanical points that facilitate large-scale conformational changes.
